# Mediodorsal thalamus contributes to the timing of instrumental actions

**DOI:** 10.1101/2020.04.18.048645

**Authors:** Nicholas Lusk, Warren H. Meck, Henry H. Yin

## Abstract

The perception of time is critical to adaptive behavior. While prefrontal cortex and basal ganglia have been implicated in interval timing in the seconds to minutes range, little is known about the role of the mediodorsal thalamus (MD), which is a key component of the limbic cortico-basal ganglia-thalamocortical loop. In this study we tested the role of the MD in timing, using an operant temporal production task in male mice. In this task, the expected timing of available rewards is indicated by lever pressing. Inactivation of the MD with muscimol produced rightward shifts in peak pressing on probe trials as well as increases in peak spread, thus significantly altering both temporal accuracy and precision. Optogenetic inhibition of glutamatergic projection neurons in the MD also resulted in similar changes in timing. The observed effects were found to be independent of significant changes in movement. Our findings suggest that the MD is a critical component of the neural circuit for interval timing, without playing a direct role in regulating ongoing performance.

**Significance Statement:** The mediodorsal nucleus of the thalamus (MD) is strongly connected with the prefrontal cortex and basal ganglia, areas which have been implicated in interval timing. Previous work has shown that the MD contributes to working memory and learning of action-outcome contingencies, but its role in behavioral timing is poorly understood. Using an operant temporal production task, we showed that inactivation of the MD significantly impaired timing behavior.

## Introduction

The ability to track the passage of time is ubiquitous across the animal kingdom and vital for survival (Buhusi and Meck, 2005). Neural correlates of timing have been found across the cortex as well as the basal ganglia (Meck, 2006; Kim et al., 2013; Yin, 2014; Finnerty et al., 2015; Gouvea et al., 2015; Bakhurin et al., 2017; Tiganj et al., 2017). Recent work has focused on the role of the prefrontal cortex (PFC) in timing (Jones et al., 2004; Buhusi et al., 2018; Kim and Narayanan, 2019). In rodents, the PFC is defined largely by the existence of strong reciprocal connections with the mediodorsal nucleus (MD) of the thalamus (Heidbreder and Groenewegen, 2003; Jones, 2012). The MD not only receives strong projections from the basal ganglia, but also sends projections that drive feed-forward inhibition within the PFC (Delevich et al., 2015). As a key thalamic component in the limbic cortico-basal ganglia-thalamocortical loop, the MD is in a position to contribute to interval timing, yet its specific functional contribution remains unclear.

Due to its strong connections with the PFC, which is often associated with working memory, considerable research has been devoted to elucidating the role of MD in working memory (Isseroff et al., 1982; Stokes and Best, 1990; Hunt and Aggleton, 1991; Watanabe and Funahashi, 2012). Although MD lesions do not reliably impair working memory(Neave et al., 1993; Hunt and Aggleton, 1998b), recent optogenetic work has demonstrated that PFC-MD interactions are crucial for working memory (Bolkan et al., 2017; Ferguson and Gao, 2018).

Previous work has also implicated the MD in the acquisition and selection of goal-directed actions controlled by action-outcome contingencies (Corbit et al., 2003; Ostlund and Balleine, 2008; Yu et al., 2012; Parnaudeau et al., 2013; Parnaudeau et al., 2015). Because animals with MD lesions fail to show sensitivity to outcome devaluation when tested on probe tests, it has been suggested that MD is critical for the representation of outcome value in guiding behavior (Alcaraz et al., 2016; Alcaraz et al., 2018). In addition, MD lesions impair acquisition and expression on a temporal differentiation task, in which animals learn to produce lever presses with specific durations to earn rewards (Yu et al., 2010). MD appears to be required when animals have to adjust the timing of their actions, consistent with the claim that the MD is critical for behavioral flexibility(Hunt and Aggleton, 1998a).

In this study, we use pharmacological and optogenetic techniques in mice to examine the role of the MD in interval timing. We first inhibited the MD during a temporal production task, using local infusions of muscimol, a GABA-A receptor agonist. MD inhibition reduced temporal precision, as indicated by peak responding on unrewarded probe trials. Optogenetic inhibition of glutamatergic projection neurons in the MD also produced similar effects on timing, resulting in rightward shifts in peak pressing when stimulation was delivered at trial onset, and lengthened a bout of lever pressing when stimulation was delivered during lever pressing.

## Materials and Methods

### Viral Constructs

AAV5-EF1α-DIO-eNpHR3.0-eYFP (halorhodopsin for optogenetic inhibition) and AAV5-EF1α-DIO-eYFP (control viral vector with no opsin) were obtained from the Duke Viral Vector Core.

### Animals

Data for muscimol experiments (n = 10) was collected from adult wild-type mice (C57BL/6J). Optogenetic experiments used adult *Vglut2-IRES-Cre* (n = 11). Male mice were used in experiments. Mice were maintained on a 12:12 light cycle. During training of 30s peak interval procedure animals were food deprived, receiving 2.0-2.5 grams of rodent chow in addition to food pellets during testing. During food deprivation animal’s weights were monitored daily to ensure they remained above 85% of their ad libitum weights. All experimental procedures were approved by the Duke University Institutional Animal Care and Use Committee.

### Surgery

Mice were anesthetized with 1.0 to 2.0% isoflurane mixed with 1.0 L/min of oxygen for surgical procedures and placed into a stereotactic frame (David Kopf Instruments, Tujunga, CA). Adult *Vglut2-IRES-Cre* mice were randomly assigned to Vglut2::eNpHR3.0 (n = 6) or Vglut2::eYFP groups (n = 5). Virus was bilaterally microinjected into the MD (0.3 μL each hemisphere, AP: −1.4 mm, ML: ± 1.48 mm, DV: −3.3 mm from skull surface, at 20° from vertical). All measurements are relative to bregma. Mice were bilaterally implanted with custom made fiber optics aimed directly above the MD (AP: −1.4 mm relative to bregma, ML: ± 1.48 mm, DV: −3.1 mm from skull surface, at 20° from vertical). Fibers were secured in place with dental acrylic adhered to skull screws. Mice were singly housed during recovery for at least two weeks before training began.

### Operant training

Lever-Press training consisted of the extension a single lever, either to the right or left of a food port. A food pellet reward (Bio-Serv 20 mg Dustless Precision Pellet) was delivered for each lever press. Every 5^th^ press lead to an alternation of which lever was extended. Sessions ended after either successfully receiving 40 rewards or 30 min. Training was considered complete after an animal received 40 rewards prior to the 30 min time limit. The median number of training sessions it took for the animals to collect 40 rewards within 30 min was 5 (2-8).

Fixed interval (FI) training sessions began with the extension of a single lever to either the left or right of the food port, counterbalanced across mice, as well as the illumination of a house light. Individual trials were demarcated by the presence of an auditory cue (white noise; 68 dB). An animal was rewarded for the first lever press occurring after the cue had been on for 30 s. At this time a food reward was delivered, the cue was turned off, and the session entered a variable inter-trial interval ranging from 90 s to 180 s randomly selected from a uniform distribution of values in 5 s increments. If there was no press within 8-seconds of the 30 s criterion, the trial would end as stated above, but without a reward. There was no penalty for presses occurring prior to 30 s. The development of a “scalloped” profile for session averaged lever pressing indicated the acquisition of timing behavior. Each session lasted 2 hrs. The median number of sessions it took to complete the FI training phase was 10 (7-27).

Peak interval (PI) training sessions consisted of two trial types: The aforementioned FI trials, and unrewarded probe trials. During probe trials, the house light was turned on for a minimum of 3× the longer target duration (90 s) plus an additional random amount of time with a mean of 20 s and a Gaussian distribution. Each trial type was randomly selected, with a 40% chance of being a probe trial. Each session lasted 150 minutes. Animals vary in how long it took for a clear peak response function to emerge on the probe trials. The median number of sessions for the PI phase was 12 (6-21) before any neural manipulations were performed.

### Local muscimol infusion

Mice received local bilateral infusions of either 0.9% saline or muscimol (0.01 mg/ml) solution. All infusions contained a total volume of 400nL (i.e. 200nL per hemisphere) infused at a rate of 0.05 mL/hr. Infusion cannulas were left in place for 10 min post infusion. After removal of cannulas, animals were again given 10 min prior to starting any training. The condition order was counterbalanced across animals. Control sessions consisted of insertion of injector cannula, but without infusing any solution. The injectors were left in place for 10 min and animals were given 10 min after removal before starting any training.

### Optogenetic stimulation

Optical stimulation began after an animal had progressed to the PI procedure and developed a distinctive peak in pressing activity centered around 30 s. Mice were connected to a 589-nm DPSS laser (Shanghai Laser) via fiber optic cables (Doric: core = 200μm; NA = 0.22) and placed inside the testing chamber (Rossi et al., 2012; Rossi et al., 2015). A rotating optical commutator (Doric) divided the beam (50:50) permitting bilateral stimulation. Stimulation (5-8 mW; constant; 15s, 10s, or 5s duration) was delivered at three timepoints: cue onset (t = 0), 15s after cue onset (t = 15), or 60s after cue onset (t = 60). The first two conditions occurred on both FI and probe trials, while the final condition only occurred on probe trials.

### Histology

After testing was completed, mice were deeply anesthetized with isoflurane and perfused with 0.1M PBS followed by 4% paraformaldehyde. Brains were post fixed in 4% paraformaldehyde for 24hrs followed by 30% sucrose solution. After sinking in the sucrose solution, brains were sliced coronally at 60 μm using a Leica CM1850 cryostat. For cannula localization, sections were mounted and immediately coverslipped with Fluoromount G with DAPI medium (Electron Microscopy Sciences; catalog #17984-24). To confirm proper fiber placement, coronal sections were compared to those from a mouse brain atlas(Paxinos and Franklin, 2003).

### Experimental design and statistical analysis

Experimental procedures were controlled by a MED Associates interface utilizing MED-PC software system. Lever presses were recorded in real time with 10 millisecond resolution. Session averaged data was analyzed using Python scripts. All statistical analysis was performed in Prism 8.

Lever pressing data for each trial was placed into 2s bins and collapsed across all trials. The cumulative pressing data was smoothed using a Savitzky-Golay filter and then fit using a gaussian curve with the addition of a linear ramp accommodating the right-tail skew, resulting from the temporal asymmetry of expected reward time relative to length of probe trials. Fits were used to obtain peak times (accuracy) and peak spread (precision) as previously described (Cheng and Meck, 2007). For optogenetic experiments fits where calculated by combining data from two sessions in order to provide a sufficient number of trials for single trial analysis, because only about half the trials of each session were stimulation trials.

Single Trial Analysis of lever pressing data was performed by putting the data in 1s bins. As the probe trials were of variable length with a minimum duration of 90s, only the first 90 bins were used for analysis. The press-rate data was padded on both sides and smoothed using a median filter, followed by a Savitzky-Golay filter. The smoothed data was log normalized and then change points were detected using python ruptures library with a Pelt search method and “L2” cost function. To account for differences in press rate between animals, the penalty and median filter length values were allowed to vary.

## Results

### Pharmacological inhibition of MD

Mice (n = 10 males) were implanted with guild cannulas prior to being trained on a 30s peak interval procedure (Figure 1A). Following experimentation, localization of cannula placement was verified for all animals (Figure 1B). After progressing to the final stage of training (see Materials & Methods) animals were assigned an infusion schedule, counterbalanced across subjects. Session averaged response rates during non-rewarded probe trials were normalized in amplitude and fit using a Gaussian + Ramp function. Control animals exhibited peak times at 26.33 ± 1.32s for the 30s PI procedure, which was lower than the 30s criteria, but in accord with previous work in mice using the same task (Buhusi et al., 2009). The peak functions revealed differences between muscimol and control conditions (Figure 1C). A repeated measures one-way ANOVA found significant effect of treatment on peak time (F(1.95, 17.58) = 10.65, p < 0.001) and variance (F(1.88, 16.90) = 20.25, p < 0.001). A Tukey’s post hoc test revealed significant increases in the muscimol peak time (33.46 ± 1.18s) and variance (15.86 ± 0.92s) relative to control (26.33 ± 1.32s, p = 0.002; 10.37 ± 0.34s, p < 0.001) as well as saline (29.19 ± 1.39s, p = 0.05; 12.07 ± 0.59s, p = 0.01). No significant differences were found between control and saline conditions for either peak time (p = 0.25) or variance (p = 0.14).

**Figure 1.**
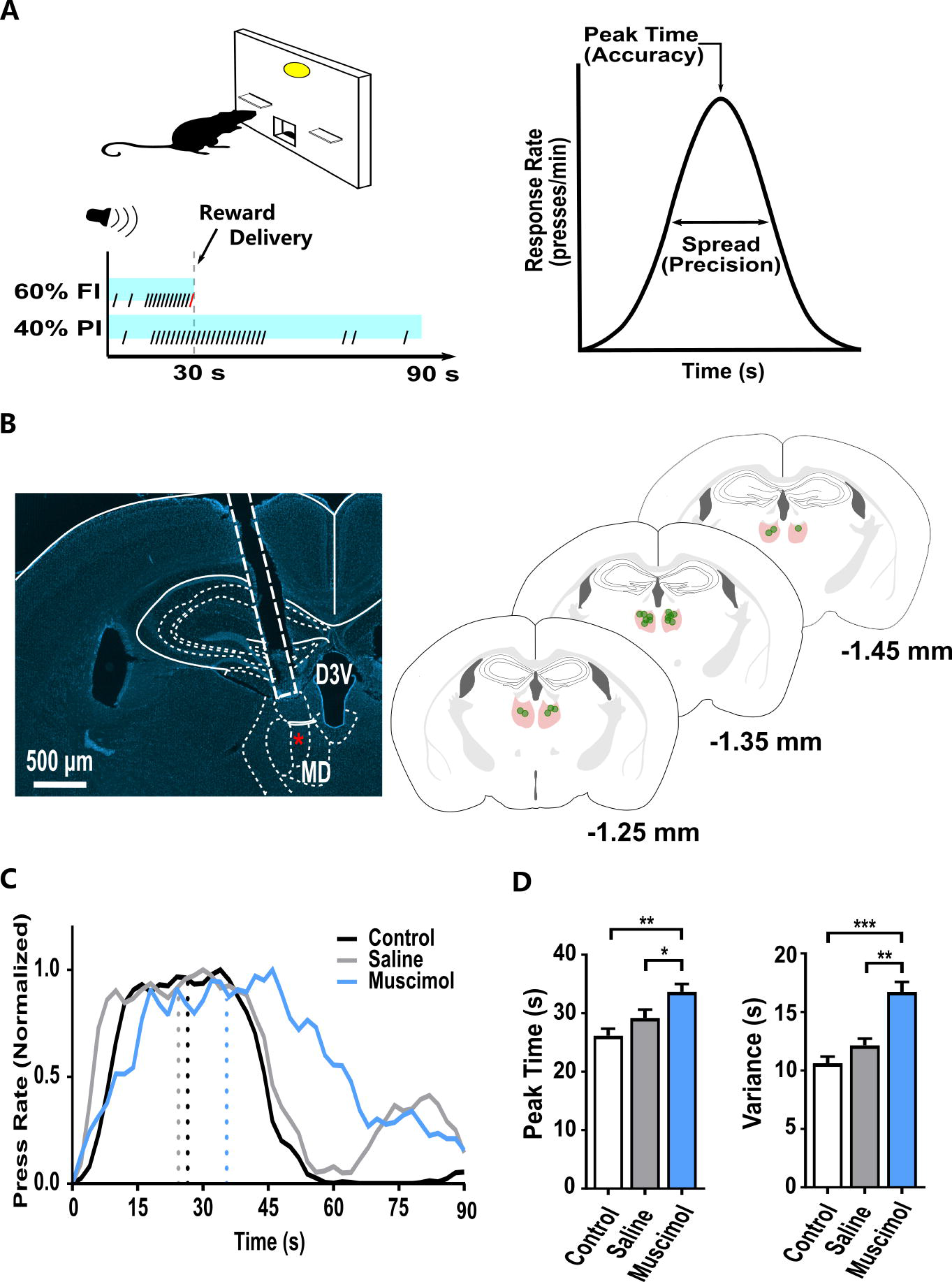
Pharmacological inhibition of MD disrupts interval timing behavior. (A) Schematic of behavioral task and idealized behavioral output. (B) Location of cannula placement. Infusion canula (red asterisk) located 0.5mm below guide canula (white box). (C) Representative performance of a mouse under control (black), saline (grey), and muscimol (blue) conditions. Dotted lines indicate peak times defined by Gaussian + Ramp fit. (D) Muscimol has effects on timing performance, from left to right: peak time, variance across, and elevation ratios. *p < 0.05, ** p < 0.01, *** p < 0.001.

To shed light on behavioral patterns that give rise to the session averaged data shown in Figure 1, we conducted a single trial analysis. For each trial, the start time, stop time, and duration of lever pressing was calculated (Figure 2A). A one-way repeated measures ANOVA found a significant treatment effect for the stop times (F(1.78, 16.02) = 24.95, p < 0.001) and duration of pressing (F(1.93, 17.41) = 14.14, p < 0.001), but not for the start times (F(1.919, 17.27) = 1.80, p = 0.20).

**Figure 2.**
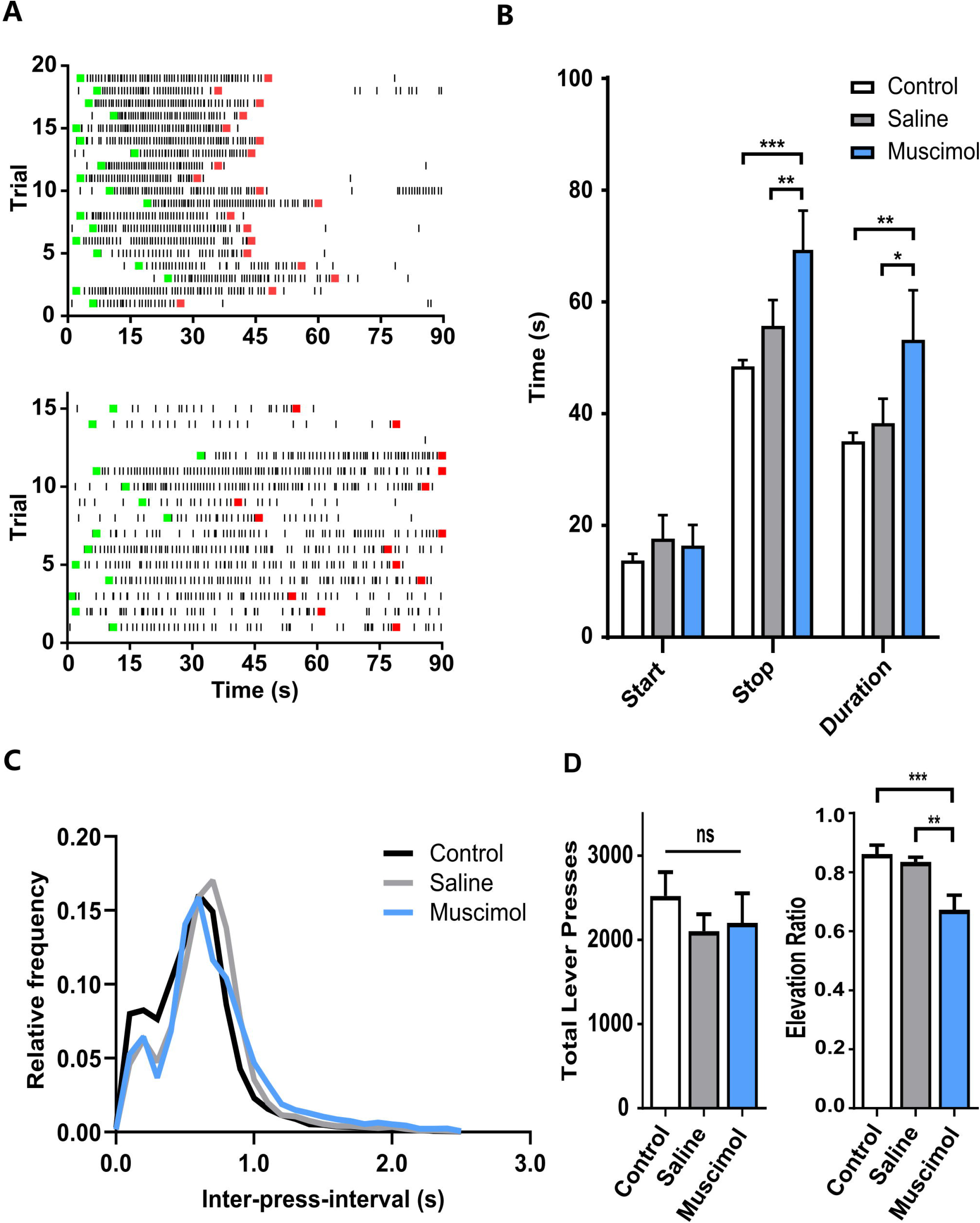
Single trial analysis demonstrates differences in pressing profiles. (A) Representative raster plot of individual trials during control (top) and muscimol (bottom) conditions from an individual mouse. Each black dash represents a single lever press. Green and red squares indicate the start and stop times, respectively (see Methods). (B) Average start time, stop time, and duration of pressing bout for mice across conditions. (C) Histogram of inter-pressintervals (IPIs) across conditions. (D) total lever presses (left) and elevation ratio (right) during a session. * p < 0.05, ** p < 0.01, *** p < 0.001

A Tukey’s post hoc test revealed significant increases in the stop time (p < 0.001; p = 0.006) as well as duration of pressing (p = 0.002; p = 0.022) for the muscimol condition compared to control and saline conditions, respectively (Figure 2B). The distribution of interpress-intervals (IPIs) shows qualitatively similar profiles between muscimol (0.69 ± 0.0044 s), saline (0.65 ± 0.0033 s) and control (0.57 ± 0.003 s) conditions (Figure 2C). Thus, the pattern of pressing is not affected by muscimol, suggesting that the observed effects cannot be due to changes in general motor activity.

In addition, no significant difference in total lever presses (F(1.98, 17.78) = 0.77, p = 0.48) nor press rate (F(1.50, 13.54) = 3.31, p = 0.078) was found, indicating increases in response variance could not be solely explained by differences in the level of motor output in muscimol animals (Figure 2D). There was also a significant main effect of treatment on elevation ratio (Buhusi et al., 2009), which measures the number of presses during the trial relative to total number presses (F(1.46, 13.12) = 21.93, p < 0.001), muscimol condition (0.70 ± 0.035) significantly lower than both control (0.87 ± 0.022, p < 0.001) and saline (0.83 ± 0.015, p = 0.007) conditions (Figure 2D). Therefore, while the muscimol doses given in this study did not significantly change overall lever pressing, it did alter sensitivity to the discriminative stimulus that indicates when presses can be rewarded.

### Optical inhibition of MD

While muscimol inhibition of the MD demonstrated its importance to proper timing behavior, this manipulation lacks temporal precision. A given trial can be divided into three phases: initial “low-state” with little pressing, followed a pressing “high-state” and concluding with another “low-state” (Church et al., 1994). To investigate the contributions of MD to behavior at different time points during the probe trial, we used optogenetics to inactivate the MD (Figure 3). We used a Cre-dependent halorhodopsin (DIO-eNpHR3.0) in vGlut2-Cre mice to selectively inhibit vGlut2+ projection neurons in the MD (Zhang et al., 2007). This manipulation provides more spatially and temporally selective inactivation of the MD. Although most MD cells are assumed to be glutamatergic, it is not possible to confirm that muscimol inactivation only affected glutamatergic neurons. By using the vGlut2-Cre driver line, stimulation can be limited to the glutamatergic projection neurons. To inactivate the MD at specific time points during the trial, we selected three timepoints for inhibition (0s, 15s, and 60s). Each timepoint occurs primarily during one of the three phases. Inhibition was constant over 15s, as a similar stimulation parameter was previously shown to effectively disrupt prefrontal dependent processes such as working memory maintenance (Bolkan et al., 2017).

**Figure 3.**
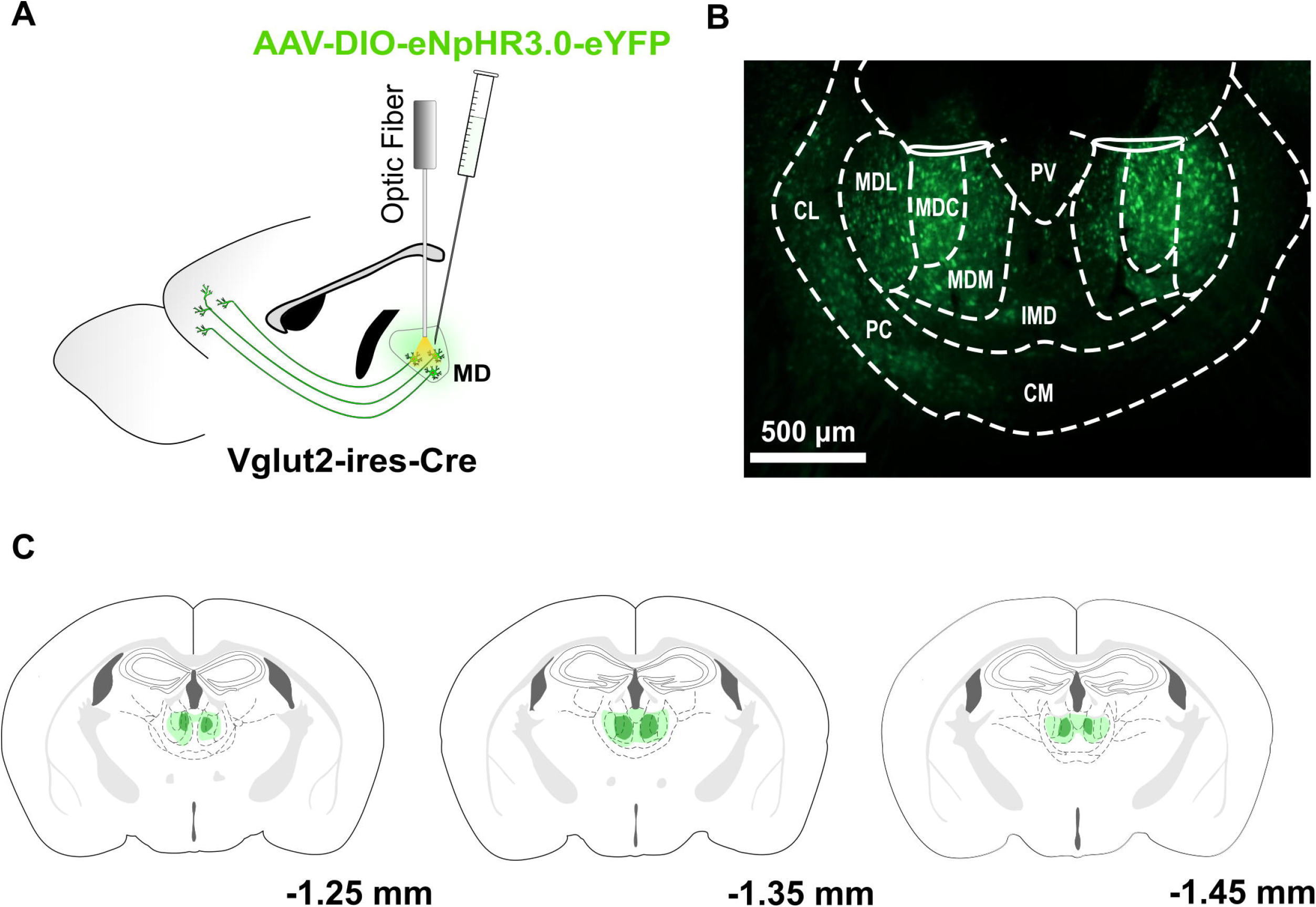
Optogenetic Experimental Design and Histology. (A) Schematic of viral infection and fiber optic implantation. (B) Histology demonstrating viral expression and fiber placement in MD. (C) Extent of viral spread indicated by the maximum (light green) and minimum (dark green) expression volume.

Performance on non-stimulation trials was consistent with previous work. Peak time (25.53s ± 0.719s) was significantly lower than 30s, but not significantly different than control data from muscimol experiments (t(14) = 0.44, p = 0.67), nor was the peak variance (t(14) = 0.84, p = 0.41). Therefore, despite having MD inhibition on 50% of the reward trials, there is likely no residual influence on non-laser peak trials. When optical inhibition was concurrent with the onset of the trial, mice showed a systematic delay in the onset of lever pressing, evidenced by a rightward shift in peak time (Figure 4A). However, the rightward shift in peak times (t(5) = 9.05, p < 0.001) was not accompanied by a significant difference in temporal variance (t(5) = 1.31, p = 0.25: Figure 4B). Single trial analysis using a multiple comparison t-test with a Holm-Sidak correction demonstrated the rightward shift to be a product of delays in the start (planned comparison, t(30) = 3.14, p = 0.011) and stop (t(30) = 2.69, p = 0.023) of the pressing high-state with no changes in the duration of pressing (t(30) = 0.45, p = 0.65: Figure 4C). A 2-dimensional density plot shows qualitative similarities in the distribution of start and stop times (Figure 4D). Despite having significantly different means, an F-test for equality of variance found no significant differences between the two conditions for start times (F(1, 26) = 2.69, p = 0.11) or stop times (F(1, 32) = 1.33, p = 0.26).

**Figure 4.**
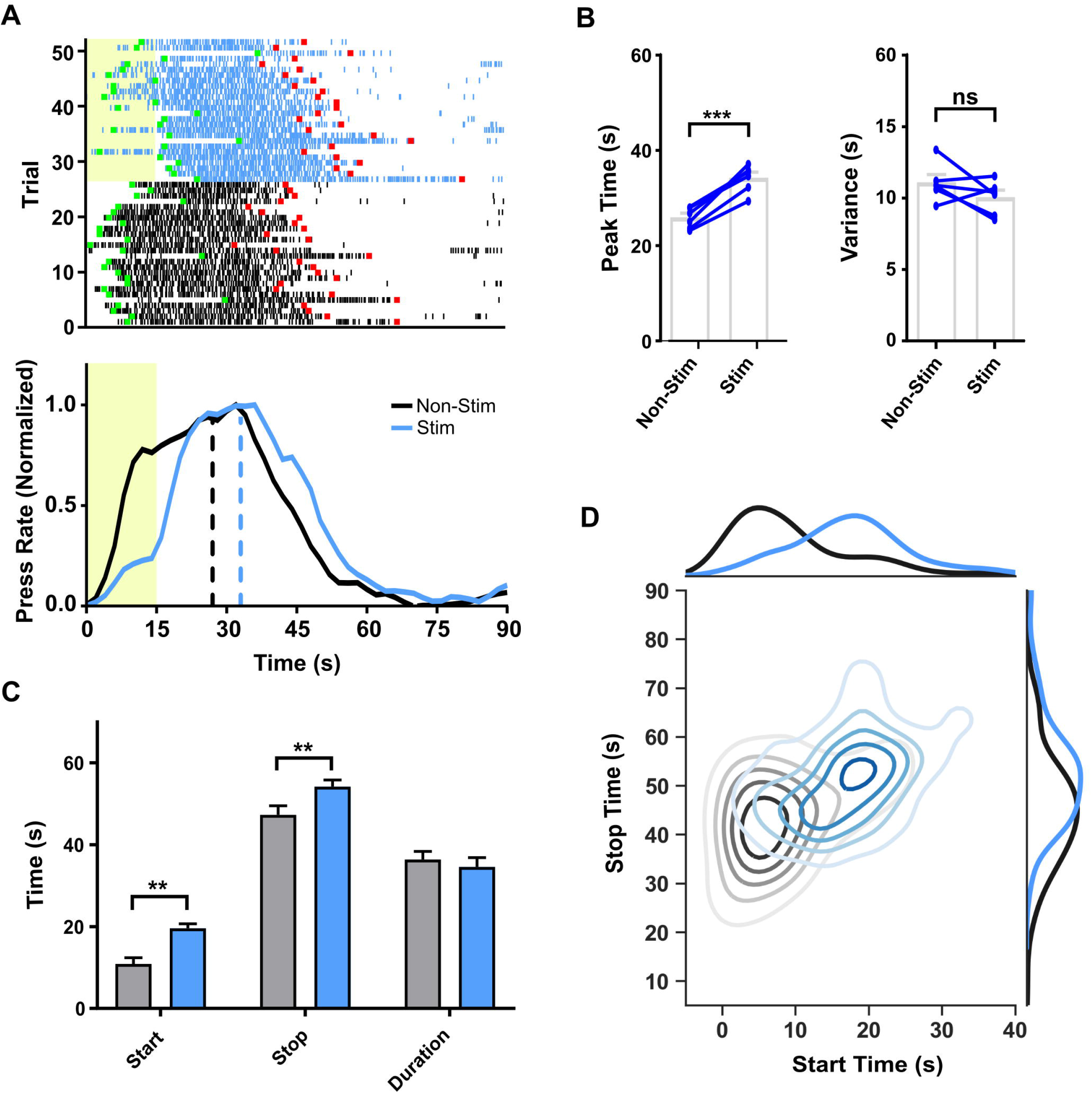
Optical inhibition of MD at cue onset delays initiation of lever pressing. (A) Schematic of viral infection and fiber optic implantation. (B) Histology demonstrating viral expression and fiber placement in MD. (C) Example of single trial (top) and averaged (bottom) pressing data over two sessions with 15s stimulation at onset of auditory cue (yellow rectangle). Each tick represents a single lever press. Green and red squares indicate the start and stop times, respectively. Dashed lines indicate peak time of Gaussian + Ramp fit. (D) Stimulation delays peak time (left) but not variance (right) of peak fits. Group average (grey) and individual (blue) data shown. (E) Stimulation significantly affects average start time and stop time, but not press bout duration. (F) Distributions of start and stop times for all trials. ** p < 0.01, *** p < 0.001

The rightward shift in peak time during optical inhibition is seemingly at odds with the findings from the muscimol experiment, which found no changes in the start time for lever pressing (Figure 2B). As mice under the muscimol condition pressed more during the inter-trial-interval (ITI), one possible explanation is that mice may already be engaged in pressing at the start of a trial. This is confirmed by analysis of lever pressing including 60 seconds (the shortest ITI duration) prior to trial onset: under muscimol more trials had start times that either preceded or coincided with the onset of the trial cue compared to the saline condition (t(9) = 2.32, p = 0.046).

Next, we started optogenetic inhibition 15s into the trial. Average start times during non-laser trials for the t = 0s condition (11.23s ± 1.001s) indicated this to be a likely time for mice to be in the early stages of their pressing. As with optical inhibition at onset, animals produce rightward shifts in peak pressing (t(5) = 5.35, p = 0.0031), but this was accompanied by a significant increase in peak variance (t(5) = 4.79, p = 0.0049: Figure 5A-B). Single trial analysis was again done in order to evaluate the contributing factors to differences found in session averaged data between conditions (Figure 5C). Contrary to the effects seen with inhibition at trial onset, inhibition at t = 15s produced significant delays in pressing stop times (t(30) = 3.596, p = 0.003) as well as increases in the duration of the pressing high-state (t(30) = 3.382, p = 0.004) with no significant differences in start times (t(15) = 0.21, p = 0.838). In addition to the increases in mean length of pressing, how inhibition of the MD affected the variability of press duration was investigated through assessment of Gaussian fits of press duration histograms (Figure 5D). An extra sum-of-squares F-test found the variance of the inhibition condition’s best fits to be significantly greater than that of non-inhibition trials (F(1, 32) = 15.14, p = 0.0005). Therefore, inhibition of the MD does not only lead to increased duration of pressing, but also increased variability in the length of a pressing bout.

**Figure 5.**
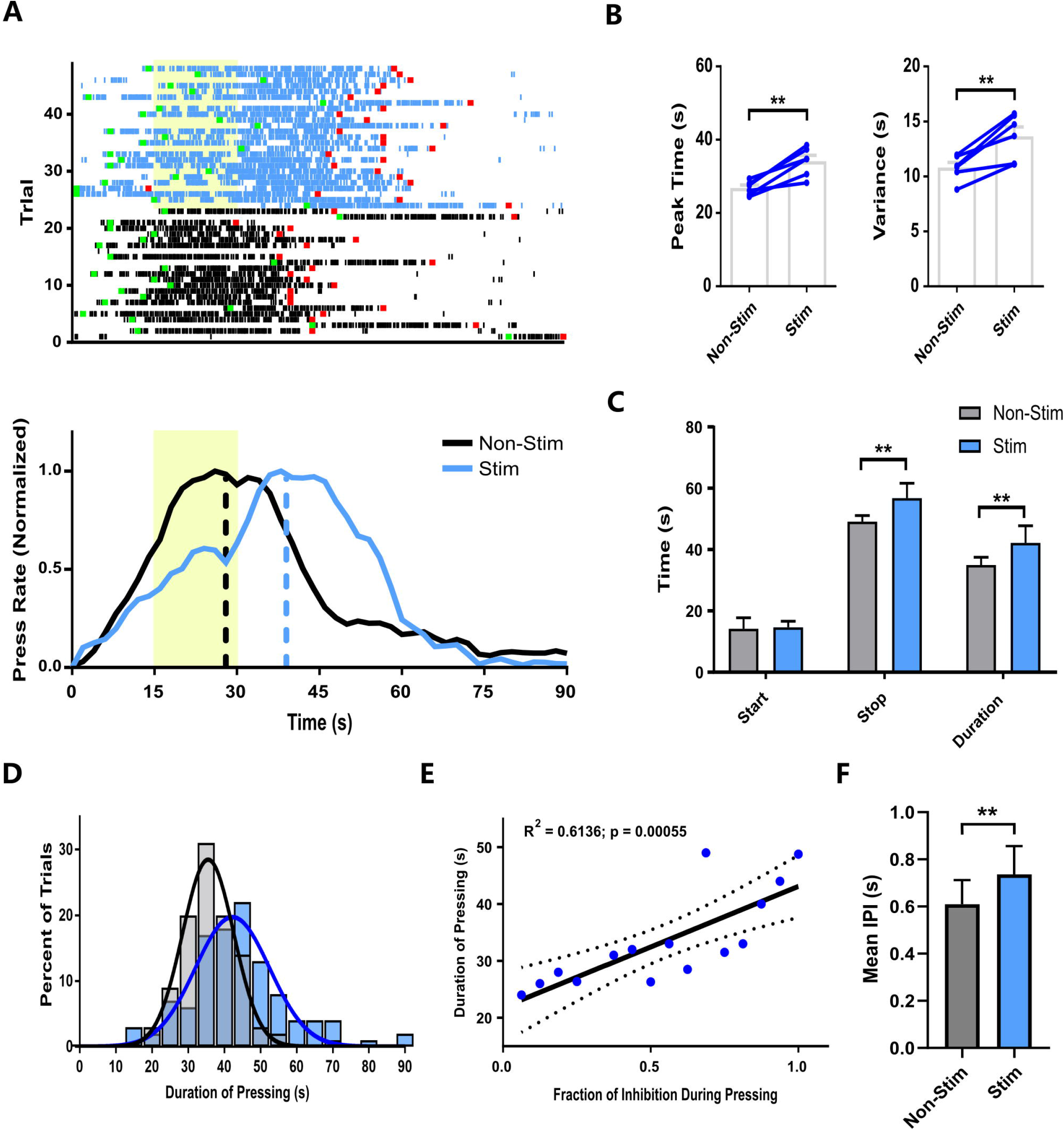
Optical inhibition of MD during pressing increases length of pressing bout. (A) Example of single trial (top) and averaged (bottom) pressing data over two sessions with 15s stimulation at onset of auditory cue (yellow rectangle). Each tick represents a single lever press. Green and red squares indicate the start and stop times, respectively. Dashed lines indicate peak time of gaussian + ramp fit (B) Stimulation delays peak time (left) and variance (right) of peak fits. Group average (grey) and individual (blue) data shown. (C) Stimulation significantly affects average start time and stop time, but not press bout duration (D) histogram of press durations for stim and non-stim trials. (E) Linear regression of press duration relative to percent of the total stim time received (F) Average inter-press-interval for each animal (n = 6) during the high-state when the laser was on (Stim) and when the laser was off (Non-Stim) *p < 0.05, ** p < 0.01

As the start of the lever pressing bout can vary from trial to trial, a linear regression was performed between the bout duration and total amount of inhibition as a fraction of the possible 15s (Figure 5E). When only part of the inhibition occurred during pressing, no distinction was made between whether inhibition occurred at the beginning or end of the pressing. The fraction of inhibition showed a strong linear correlation with duration of the lever pressing bout (R^2^ = 0.61, p = 0.00055). To test whether optical inhibition reduces the rate of pressing, the interpress-interval (IPI) within the high-state during the time of optical inhibition was compared to the IPI within the high-state when optical inhibition was not occurring (Figure 5F). Surprisingly, a paired t-test found that during optical inhibition mice pressed more rapidly, evidenced by a significantly shorter IPI (t(5) = −4.05, p = 0.0098). While this finding demonstrates a degree of hyperactivity, it rules out the possibility that a slowing of motor output could explain the timing effects.

The final time point for optical inhibition, t = 60s, occurs after the animal has finished pressing. As expected, inhibition at this point has no significant effects on performance (session averaged, Figures 6A-B; single trial, Figures 6C). However, mice exhibited a significant increase in rebound pressing during inhibition trials (Figure 6D: t(5) = 3.30, p = 0.030). This is unlikely to be explained by confounding factors such as visually perceiving light from the laser as eYFP-control mice did not exhibit any changes in rebound pressing (t(5) = 0.031, p = 0.49).

**Figure 6.**
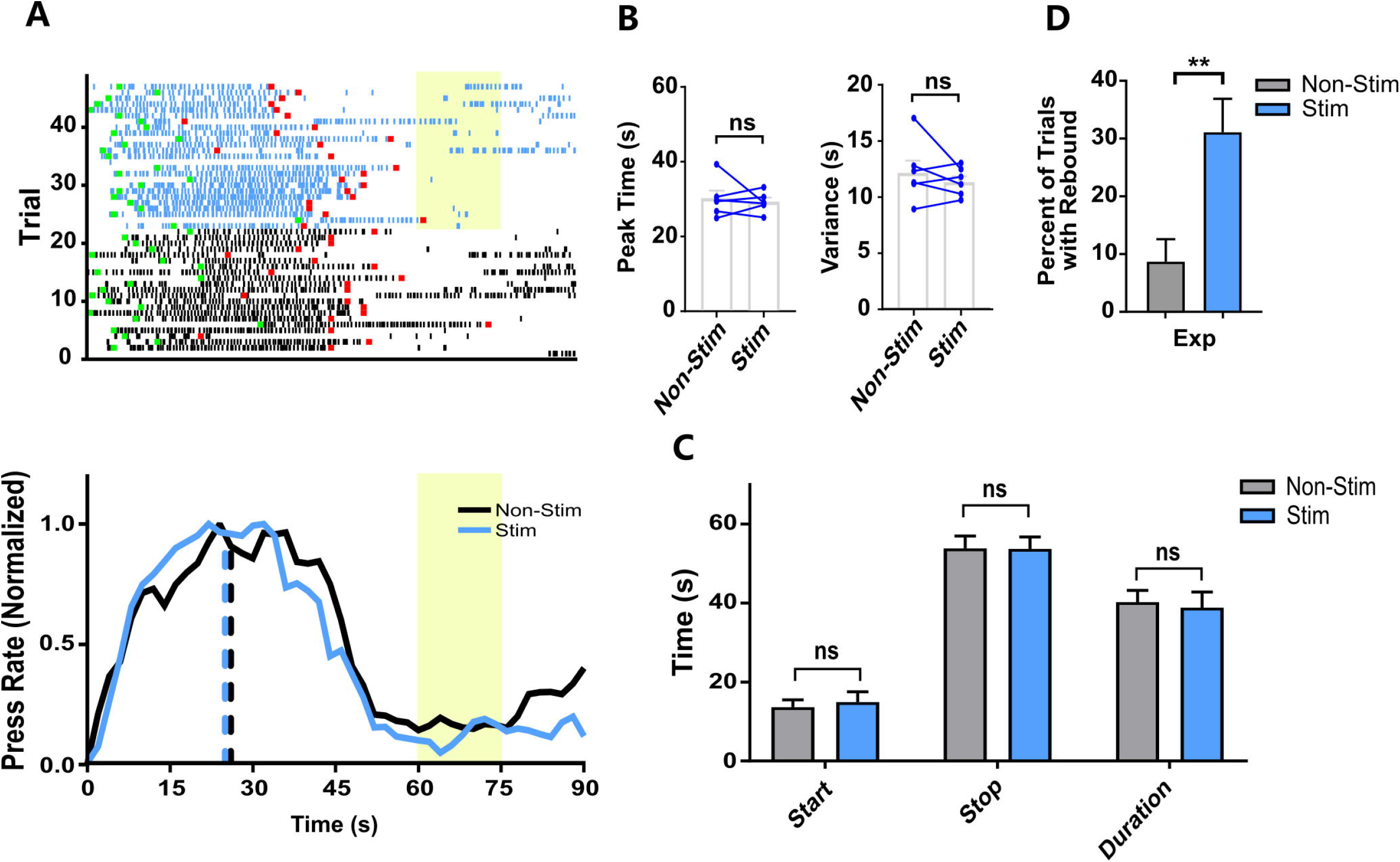
Optical inhibition following pressing increases probability of reengagement. (A) Example of single trial (top) and averaged (bottom) pressing data over two sessions with 15s stimulation at t = 60s of auditory cue (yellow rectangle). Each tick represents a single lever press. Green and red squares indicate the start and stop times, respectively. Dashed lines indicate peak time of gaussian + ramp fit. (B) Stimulation has no effect on peak time (left) or variance (right) of peak fits. Group average (grey) and individual (blue) data shown. (C) Stimulation has no significant affect start time and stop time, or press bout duration. (D) Percent of trials with reengagement (rebound). ** p < 0.01

To address potential confounds related to light delivery, AAV5-EF1α-DIO-eYFP controls (n = 5) were tested on the 30s PI procedure using identical parameters. A paired t-test on session averaged data showed no difference in peak times (p = 0.24, p = 0.20, p = 0.090) nor variance (p = 0.43, p = 0.091, p = 0.84) for control mice at stimulation times t = 0s, 15s, and 60s, respectively (Figure 7 A-C). Additionally, single trial analysis at all three stimulation timepoints was conducted to confirm averaging effects did not mask differences at a lower level of analysis. There were no differences in start (p = 0.74, p = 0.89, p = 0.57), stop (p = 0.94, p = 0.80, p = 0.64) or duration (p = 0.75, p = 0.72, p = 0.84), respectively (Figure 7D).

**Figure 7.**
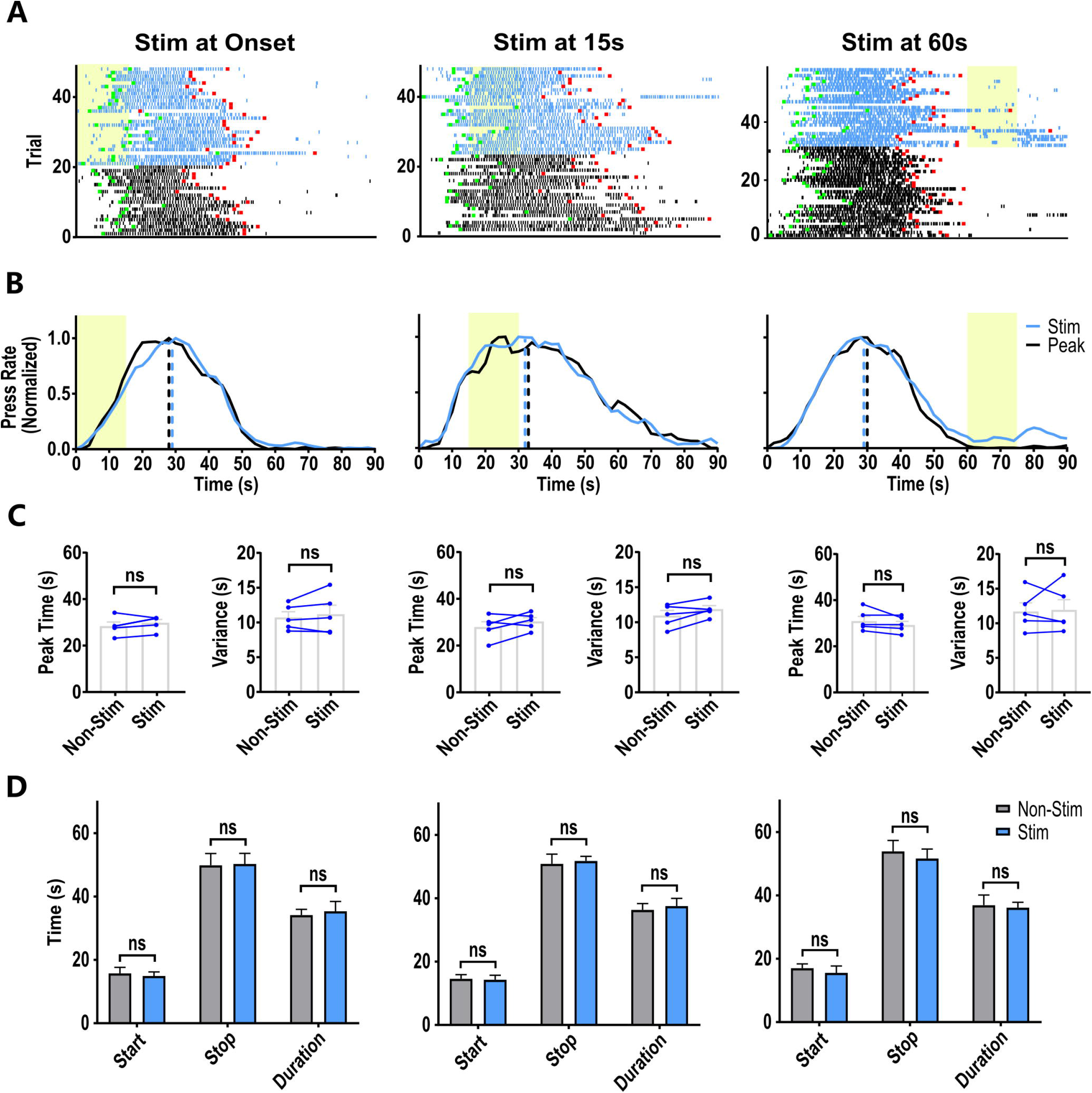
Photo-stimulation did not change behavior in eYFP controls. (A) Raster plots for eYFP control mice for at timepoints, from left to right, t = 0s, t = 15s, and t = 60s. (B) Session averaged data for eYFP control animals for stim (blue) and non-stim (black) trials at timepoints, from left to right, t = 0s, t = 15s, and t = 60s (yellow). Peak times for stim (blue dash) and non-stim (black dash) defined by Gaussian + Ramp fits. (C) Stimulation does not modify peak time (left) or variance (right) of peak fits. Group average (grey) and individual (blue) data shown. (D) Stimulation has no significant affect average start time and stop time, or press bout duration.

Lastly, we investigated how the length of optical inhibition modulates timing behavior at all three time points across varies lengths of inhibition (5s, 10s and 15s). For optical inhibition at trial onset (t = 0s), a repeated measures ANOVA found a significant main effect of stimulation length (F(2, 8) = 9.85, p = 0.007). Post hoc tests found a significant difference between 15s of inhibition and 5s (t(8) = 4.29, p = 0.0079) as well as 10s (t(8) = 3.13, p = 0.042: Figure 8A). Moreover, inhibition at t = 15s demonstrated a significant treatment effect (F(2, 8) = 30.21, p < 0.001) with longer durations of MD inhibition leading to increases in variance. A post-hoc test found 15s inhibition to produce significantly greater increases in variance than 5s or 10s of inhibition (t(8) = 7.39, p < 0.001; t(8) = 5.77, p = 0.001), respectively (Figure 8B). Moreover, this finding aligns with single trial data demonstrating increases in the length of the high-state correlate with the length of optical inhibition that occurs over the pressing phase. A two-way repeated measures ANOVA with stimulation duration (5, 10, 15s) and stimulation condition (stimulation, non-stimulation) as factors revealed a significant main effect of duration on percentage of rebound trials (F(2, 8) = 5.20, p = 0.036), but no main effect of stimulation condition or interaction between these factors. Post hoc tests found inhibition lengths of 15s (t(8) = 5.53, p = 0.002) and 10s (t(8) = 4.44, p = 0.007) result in a higher proportion of rebound trials than non-inhibition trials (Figure 8C). Additionally, the 5s inhibition condition did not significantly increase rebound pressing (t(8) = 1.56, p = 0.47).

**Figure 8.**
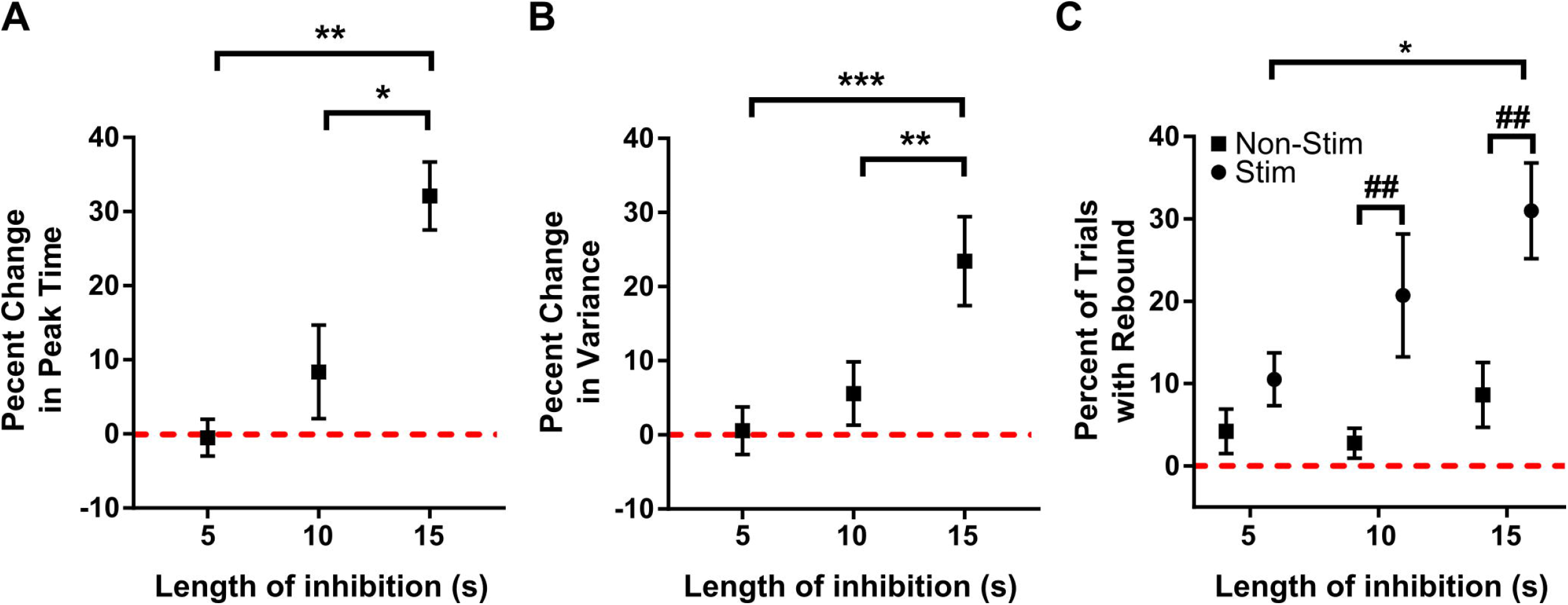
Effects of optical inhibition scale with length of inhibition. (A) Percent increase in session peak time relative to length of optical inhibition at t = 0s. (B) Percent increase in session variance relative to length of optical inhibition at t = 15s. (C) Percent of trial with rebound pressing relative to length of optical inhibition at t = 60s. Within condition * p < 0.05, ** p < 0.01, *** p < 0.001, across condition ## p < 0.01.

## Discussion

Utilizing pharmacological and optogenetic manipulations, we provide evidence for the involvement of the MD in interval timing. We first trained mice on an FI task, in which rewards become available after 30 seconds. The internal estimate of time is indicated by the peak lever pressing on occasional probe trials with no reward. Here we show rightward shifts in the peak timing function in addition to increases in timing variance after MD inactivation. Such effects are not confounded by general changes in motor output, and therefore likely due to changes in internal timing mechanisms.

Reversible pharmacological inhibition of the MD with muscimol modified timing performance, impairing both timing accuracy and precision, as indicated by peak time and spread of session averaged performance on probe trials. In addition to timing behavior, mice exhibited a decrease in the ratio of presses during cued trials quantified through the elevation ratio (Figure 1–2). These findings resemble the effects of mPFC inactivation as muscimol inactivation of the prelimbic (PL) cortex also produced similar effects (Buhusi et al., 2018). Additionally, mixed effects on response rate were found depending on dosage of muscimol with low doses leading to increases in response rate. This is akin to the discrepancy found in the current work with the muscimol dosage used producing no significant changes in response rate, but the optogenetic inhibition leading to increases in response rate. The strong reciprocal connectivity between the MD and prefrontal cortical areas and similarities in effects on timing from inactivation of these two regions suggests a critical role of cortico-thalamo-cortical interactions in interval timing.

To further elucidate MD contribution to various stages of timing behavior, we used optogenetics to inhibit MD output neurons at various time points. Inhibition of the MD was found to increase the duration of the animal’s lever pressing bout during a trial, resulting in delays in the onset of pressing (Figure 4) and offset of pressing (Figure 5). Duration of inhibition during pressing strongly correlates with the increase in the length of the pressing bout. This suggests that the mouse may time how long to wait before starting to press and then time how long to press once initiated. That stimulation duration during pressing correlates with the length of the pressing bout suggests each stage of the task is timed independently. That is, the animal will time when it should begin pressing and then when to stop, sequentially. This interpretation of timing was proposed previously (Killeen and Fetterman, 1988). However, this is in conflict with early analysis of individual trial studies that found support for parallel processing theories, most notably the Scalar Expectancy Theory, according to which all behavioral output is generated in relation to a single timed duration (Gibbon, 1977). Previous work found the covariance pattern between start and stop times of pressing bouts supported the use of a single temporal sample from memory with different decision thresholds for when to start and stop pressing(Church et al., 1994). However, the ability to maintain the necessary covariance pattern does not require parallel processing, as theories based on memory decay have demonstrated (Staddon and Higa, 1999). Our results favor a serial timing process in which animals time the length of each individual stage during a peak interval procedure.

Our findings also have implications for computational models of interval timing. Many timing models focus on short intervals in the sub-second range, but neural networks relying solely on cortical based connectivity rules have trouble scaling to longer time intervals (>10 s) (Laje and Buonomano, 2013). It has been suggested that shifts in the balance of excitation and inhibition (E/I) within thalamocortical networks allow for the emergence and stabilization of sequential activity patterns(Hardy and Buonomano, 2016). Recent work has suggested that the MD is important for maintaining E/I balance within the mPFC (Ferguson and Gao, 2018), by modulating feedforward inhibition via GABAergic interneurons in the cortex (Delevich et al., 2015). Adjusting E/I balance via cortico-thalamo-cortical interactions may improve the maintenance of working memory in these networks and allow timing of longer durations.

Thus we show for the first time the importance of the MD in timing long intervals. The durations used in the current work span 30 seconds, much long than that used in previous work on the role of thalamus in timing (Wang et al., 2018). While shorter durations (i.e. < 2s) can be effectively timed in localized networks such as the cerebellum (Johansson et al., 2014), as the relevant time interval increases, neural substrate of timing appears to require a larger network of brain regions, requiring synchronization, coordination, or integration of temporal information from the distributed sources (Matell and Meck, 2004; Hass and Durstewitz, 2016; Petter et al., 2016). Substantial evidence supports the role of higher-order thalamic nuclei, such as the MD, in synchronizing neural activity in the cortex (Theyel et al., 2010; Saalmann, 2014). This role of the MD also explains why changes in timing behavior after MD inactivation resemble those found after inactivation of connected cortical regions(Buhusi et al., 2018).

In addition to interactions with the frontal cortex, the MD is also a major target of basal ganglia output nuclei such as the substantia nigra pars reticulata (SNr)(Gerfen et al., 1982; Haber et al., 1985; Kuroda and Price, 1991; Miyamoto and Jinnai, 1994). Extensive work has established the role of the basal ganglia circuit in interval timing (Matell and Meck, 2004; Gouvea et al., 2015; Bakhurin et al., 2020). However, recent optogenetic work also has implicated the nigral output in both ongoing performance and timing of future behavior. While the nigral output is known to play a key role in posture control and top-down regulation of movement parameters(Barter et al., 2014; Barter et al., 2015), activation of the GABAergic nigrotectal projection not only reduces ongoing licking but proportionally delays subsequently timed behavior (Toda et al., 2017). However, in the present study, MD inhibition does not appear to impair ongoing performance. It is possible that manipulations of the SNr can alter performance via direct projections to the tectum and brainstem pattern generators (Rossi et al., 2016). However, collaterals of nigral efferent projections may send a copy of the ongoing motor commands to the MD. This is consistent with previous work in monkeys showing that the MD does not play a role in movement generation, but is critical for internal monitoring of movement sequences by signaling an efference copy or corollary discharge to frontal cortical regions (Sommer and Wurtz, 2002). Although projections from the SNr usually target more lateral regions of the MD than those targeted in the current work, it is possible that there is significant overlap in the target region and the region inactivated in our experiments, or that different regions in the MD may signal different types of efference copy signals corresponding to different types of actions. Future work targeting specific SNr-MD and MD-PFC pathways may shed light on how efference copy signals are computed and used in the timing of instrumental actions.

## Acknowledgments

We would like to dedicate this paper to the memory of our co-author Warren H. Meck, a pioneer in the study of interval timing, who passed away unexpectedly during the preparation of the manuscript. This research was supported by DA040701, MH112883, and NS094754 to HHY.

## References

Alcaraz F, Naneix F, Desfosses E, Marchand AR, Wolff M, Coutureau E (2016) Dissociable effects of anterior and mediodorsal thalamic lesions on spatial goal-directed behavior. Brain Structure and Function 221:79–89.

Alcaraz F, Fresno V, Marchand AR, Kremer EJ, Coutureau E, Wolff M (2018) Thalamocortical and corticothalamic pathways differentially contribute to goal-directed behaviors in the rat. eLife 7:e32517.

Bakhurin KI, Goudar V, Shobe JL, Claar LD, Buonomano DV, Masmanidis SC (2017) Differential Encoding of Time by Prefrontal and Striatal Network Dynamics. The Journal of neuroscience: the official journal of the Society for Neuroscience 37:854–870.

Bakhurin KI, Li X, Friedman AD, Lusk NA, Watson GD, Kim N, Yin HH (2020) Opponent regulation of action performance and timing by striatonigral and striatopallidal pathways. eLife 9:e54831.

Barter JW, Castro S, Sukharnikova T, Rossi MA, Yin HH (2014) The role of the substantia nigra in posture control. European Journal of Neuroscience 39 (9):1465–1473.

Barter JW, Li S, Sukharnikova T, Rossi MA, Bartholomew RA, Yin HH (2015) Basal ganglia outputs map instantaneous position coordinates during behavior. Journal of Neuroscience 35:2703–2716.

Bolkan SS, Stujenske JM, Parnaudeau S, Spellman TJ, Rauffenbart C, Abbas AI, Harris AZ, Gordon JA, Kellendonk C (2017) Thalamic projections sustain prefrontal activity during working memory maintenance. Nature neuroscience 20:987.

Buhusi CV, Meck WH (2005) What makes us tick? Functional and neural mechanisms of interval timing. Nature reviews Neuroscience.

Buhusi CV, Reyes MB, Gathers C-A, Oprisan SA, Buhusi M (2018) Inactivation of the medial-prefrontal cortex impairs interval timing precision, but not timing accuracy or scalar timing in a peak-interval procedure in rats. Frontiers in Integrative Neuroscience 12:20.

Buhusi CV, Aziz D, Winslow D, Carter RE, Swearingen JE, Buhusi MC (2009) Interval timing accuracy and scalar timing in C57BL/6 mice. Behav Neurosci 123:1102–1113.

Cheng R-K, Meck WH (2007) Prenatal choline supplementation increases sensitivity to time by reducing non-scalar sources of variance in adult temporal processing. Brain research 1186:242–254.

Church RM, Meck WH, Gibbon J (1994) Application of scalar timing theory to individual trials. Journal of Experimental Psychology: Animal Behavior Processes 20:135.

Corbit LH, Muir JL, Balleine BW (2003) Lesions of mediodorsal thalamus and anterior thalamic nuclei produce dissociable effects on instrumental conditioning in rats. The European journal of neuroscience 18:1286–1294.

Delevich K, Tucciarone J, Huang ZJ, Li B (2015) The mediodorsal thalamus drives feedforward inhibition in the anterior cingulate cortex via parvalbumin interneurons. Journal of Neuroscience 35:5743–5753.

Ferguson BR, Gao W-J (2018) Thalamic control of cognition and social behavior via regulation of gamma-aminobutyric acidergic signaling and excitation/inhibition balance in the medial prefrontal cortex. Biological psychiatry 83:657–669.

Finnerty GT, Shadlen MN, Jazayeri M, Nobre AC, Buonomano DV (2015) Time in cortical circuits. Journal of Neuroscience 35:13912–13916.

Gerfen C, Staines WA, Fibiger HC, Arbuthnott GW (1982) Crossed connections of the substantia nigra in the rat. Journal of Comparative Neurology 207:283–303.

Gibbon J (1977) Scalar Expectancy-Theory and Webers Law in Animal Timing. Psychological Review 84:279–325.

Gouvea TS, Monteiro T, Motiwala A, Soares S, Machens C, Paton JJ (2015) Striatal dynamics explain duration judgments. eLife 4.

Haber SN, Groenewegen HJ, Grove EA, Nauta WJ (1985) Efferent connections of the ventral pallidum: evidence of a dual striato pallidofugal pathway. The Journal of comparative neurology 235:322–335.

Hardy NF, Buonomano DV (2016) Neurocomputational models of interval and pattern timing. Current Opinion in Behavioral Sciences 8:250–257.

Hass J, Durstewitz D (2016) Time at the center, or time at the side? Assessing current models of time perception. Current Opinion in Behavioral Sciences 8:238–244.

Heidbreder CA, Groenewegen HJ (2003) The medial prefrontal cortex in the rat: evidence for a dorso-ventral distinction based upon functional and anatomical characteristics. Neuroscience & Biobehavioral Reviews 27:555–579.

Hunt PR, Aggleton JP (1991) Medial dorsal thalamic lesions and working memory in the rat. Behavioral and neural biology 55:227–246.

Hunt PR, Aggleton JP (1998a) Neurotoxic lesions of the dorsomedial thalamus impair the acquisition but not the performance of delayed matching to place by rats: a deficit in shifting response rules. Journal of Neuroscience 18:10045–10052.

Hunt PR, Aggleton JP (1998b) An examination of the spatial working memory deficit following neurotoxic medial dorsal thalamic lesions in rats. Behavioural brain research 97:129–141.

Isseroff A, Rosvold H, Galkin T, Goldman-Rakic P (1982) Spatial memory impairments following damage to the mediodorsal nucleus of the thalamus in rhesus monkeys. Brain research 232:97–113.

Johansson F, Jirenhed D-A, Rasmussen A, Zucca R, Hesslow G (2014) Memory trace and timing mechanism localized to cerebellar Purkinje cells. Proceedings of the National Academy of Sciences 111:14930–14934.

Jones CR, Rosenkranz K, Rothwell JC, Jahanshahi M (2004) The right dorsolateral prefrontal cortex is essential in time reproduction: an investigation with repetitive transcranial magnetic stimulation. Experimental Brain Research 158:366–372.

Jones EG (2012) The thalamus: Springer Science & Business Media.

Killeen PR, Fetterman JG (1988) A behavioral theory of timing. Psychological review 95:274.

Kim J, Ghim J-W, Lee JH, Jung MW (2013) Neural correlates of interval timing in rodent prefrontal cortex. Journal of Neuroscience 33:13834–13847.

Kim Y-C, Narayanan NS (2019) Prefrontal D1 dopamine-receptor neurons and delta resonance in interval timing. Cerebral Cortex 29:2051–2060.

Kuroda M, Price JL (1991) Ultrastructure and synaptic organization of axon terminals from brainstem structures to the mediodorsal thalamic nucleus of the rat. Journal of Comparative Neurology 313:539–552.

Laje R, Buonomano DV (2013) Robust timing and motor patterns by taming chaos in recurrent neural networks. Nature neuroscience 16:925.

Matell MS, Meck WH (2004) Cortico-striatal circuits and interval timing: coincidence detection of oscillatory processes. Cognitive brain research 21:139–170.

Meck WH (2006) Neuroanatomical localization of an internal clock: a functional link between mesolimbic, nigrostriatal, and mesocortical dopaminergic systems. Brain Res 1109:93–107.

Miyamoto Y, Jinnai K (1994) The inhibitory input from the substantia nigra to the mediodorsal nucleus neurons projecting to the prefrontal cortex in the cat. Brain research 649:313–318.

Neave N, Sahgal A, Aggleton JP (1993) Lack of effect of dorsomedial thalamic lesions on automated tests of spatial memory in the rat. Behavioural brain research 55:39–49.

Ostlund SB, Balleine BW (2008) Differential involvement of the basolateral amygdala and mediodorsal thalamus in instrumental action selection. The Journal of neuroscience: the official journal of the Society for Neuroscience 28:4398–4405.

Parnaudeau S, Taylor K, Bolkan SS, Ward RD, Balsam PD, Kellendonk C (2015) Mediodorsal thalamus hypofunction impairs flexible goal-directed behavior. Biological psychiatry 77:445–453.

Parnaudeau S, O’Neill P-K, Bolkan SS, Ward RD, Abbas AI, Roth BL, Balsam PD, Gordon JA, Kellendonk C (2013) Inhibition of mediodorsal thalamus disrupts thalamofrontal connectivity and cognition. Neuron 77:1151–1162.

Paxinos G, Franklin K (2003) The mouse brain in stereotaxic coordinates. New York: Academic Press.

Petter EA, Lusk NA, Hesslow G, Meck WH (2016) Interactive roles of the cerebellum and striatum in sub-second and supra-second timing: Support for an initiation, continuation, adjustment, and termination (ICAT) model of temporal processing. Neuroscience & Biobehavioral Reviews 71:739–755.

Rossi MA, Calakos N, Yin HH (2015) Spotlight on movement disorders: what optogenetics has to offer. Movement Disorders 30:624–631.

Rossi MA, Hayrapetyan VY, Maimon B, Mak K, Je HS, Yin HH (2012) Prefrontal cortical mechanisms underlying delayed alternation in mice. Journal of neurophysiology 108:1211–1222.

Rossi MA, Li HE, Lu D, Kim IH, Bartholomew RA, Gaidis E, Barter JW, Kim N, Cai MT, Soderling SH, Yin HH (2016) A GABAergic nigrotectal pathway for coordination of drinking behavior. Nature neuroscience 19:742–748.

Saalmann YB (2014) Intralaminar and medial thalamic influence on cortical synchrony, information transmission and cognition. Frontiers in systems neuroscience 8:83.

Sommer MA, Wurtz RH (2002) A pathway in primate brain for internal monitoring of movements. Science 296:1480–1482.

Staddon J, Higa J (1999) Time and memory: Towards a pacemaker-free theory of interval timing. Journal of the experimental analysis of behavior 71:215–251.

Stokes KA, Best PJ (1990) Mediodorsal thalamic lesions impair “reference” and “working” memory in rats. Physiology & behavior 47:471–476.

Theyel BB, Llano DA, Sherman SM (2010) The corticothalamocortical circuit drives higher-order cortex in the mouse. Nature neuroscience 13:84–88.

Tiganj Z, Jung MW, Kim J, Howard MW (2017) Sequential firing codes for time in rodent medial prefrontal cortex. Cerebral Cortex 27:5663–5671.

Toda K, Lusk NA, Watson GDR, Kim N, Lu D, Li HE, Meck WH, Yin HH (2017) Nigrotectal Stimulation Stops Interval Timing in Mice. Current biology: CB 27:3763–3770 e3763.

Wang J, Narain D, Hosseini EA, Jazayeri M (2018) Flexible timing by temporal scaling of cortical responses. Nature neuroscience 21:102–110.

Watanabe Y, Funahashi S (2012) Thalamic mediodorsal nucleus and working memory. Neuroscience & Biobehavioral Reviews 36:134–142.

Yin HH (2014) Action, time and the basal ganglia. Philosophical Transactions of the Royal Society B: Biological Sciences 369.

Yu C, Gupta J, Yin HH (2010) The role of mediodorsal thalamus in temporal differentiation of reward-guided actions. Front Integr Neurosci 4.

Yu C, Fan D, Lopez A, Yin HH (2012) Dynamic changes in single unit activity and gamma oscillations in a thalamocortical circuit during rapid instrumental learning. PLoS One 7:e50578.

Zhang F, Wang L-P, Brauner M, Liewald JF, Kay K, Watzke N, Wood PG, Bamberg E, Nagel G, Gottschalk A (2007) Multimodal fast optical interrogation of neural circuitry. Nature 446:633–639.

